# Stop codon recoding is widespread in diverse phage lineages and has the potential to regulate translation of late stage and lytic genes

**DOI:** 10.1101/2021.08.26.457843

**Authors:** Adair L. Borges, Yue Clare Lou, Rohan Sachdeva, Basem Al-Shayeb, Alexander L. Jaffe, Shufei Lei, Joanne M. Santini, Jillian F. Banfield

## Abstract

The genetic code is a highly conserved feature of life. However, some “alternative” genetic codes use reassigned stop codons to code for amino acids. Here, we survey stop codon recoding across bacteriophages (phages) in human and animal gut microbiomes. We find that stop codon recoding has evolved in diverse clades of phages predicted to infect hosts that use the standard code. We provide evidence for an evolutionary path towards recoding involving reduction in the frequency of TGA and TAG stop codons due to low GC content, followed by acquisition of suppressor tRNAs and the emergence of recoded stop codons in structural and lysis genes. In analyses of two distinct lineages of recoded virulent phages, we find that lysis-related genes are uniquely biased towards use of recoded stop codons. This convergence supports the inference that stop codon recoding is a strategy to regulate the expression of late stage genes and control lysis timing. Interestingly, we identified prophages with recoded stop codons integrated into genomes of bacteria that use standard code, and hypothesize that recoding may control the lytic-lysogenic switch. Alternative coding has evolved many times, often in closely related lineages, indicating that genetic code is plastic in bacteriophages and adaptive recoding can occur over very short evolutionary timescales.

## Introduction

The genetic code is highly conserved and largely considered to be invariant and evolutionarily static^1^. However, a subset of organisms employ “alternative” genetic codes, which involves the reassignment of one or more codons^2^. In eukaryotes, alternatively coded genomes include the nuclear genomes of some ciliates^3–5^, diplomonads^6^, green algae^7^, and yeasts^8,9^. Many endosymbiont genomes are recoded^2^, including the vertebrate mitochondrial genome^10^. In prokaryotes, Mycoplasma^11,12^ and Spiroplasma^13^ have recoded the TGA stop codon to tryptophan (W), and members of the Candidate Phyla Radiation (CPR) Gracilibacteria and Absconditabacteria have recoded the TGA stop codon to glycine^14–16^ (G). Fascinatingly, some large uncultivated phages of the gut microbiome (predominantly Lak phages and crAssphages) have recoded the TAG or TGA stop codon^17–21^. Importantly, alternatively coded Lak^18,21^ and crAssphage^20,22^ are predicted to infect members of the *Bacteroidetes* phylum, which use standard code. Thus, alternatively coded phages represent a unique situation: unlike other recoded organisms which changed their translation system on evolutionary timescales, alternatively coded phages must convert the translation environment of their infected host cell from standard code to alternative code in a matter of minutes.

It is unknown why phages have evolved a biological feature as fundamentally incompatible with their host translation system as a disparate genetic code. An analysis of a single alternatively coded phage genome^17^ (now understood to be a crAssphage^20^) hypothesized that alternative coding could be a manifestation of phage-host antagonism, though this argument has not been explored further. In this model, the TAG-recoded crAssphage was hypothesized to infect TGA-recoded bacteria, where phage and host would mutually disrupt translation of the others’ genes via release factor activity. Furthermore, it is unknown if alternative coding in large phages such as crAssphage and Lak phage represents an unusual characteristic of some classes of large phages or if it is a more widespread feature of phage populations.

Here, we survey stop codon recoding in bacteriophages of the human and animal microbiome. We identify diverse lineages of phages with recoded stop codons that are predicted to infect bacteria that use standard code, and show that code change can happen over very short evolutionary distances. We infer that stop codon recoding is a mode of post-transcriptional regulation used to control critical points in the phage life cycle, such as the establishment of lysogeny and the triggering of cell lysis.

## Results

### Incidence of bacteriophage stop codon recoding in human and animal microbiomes

We systematically evaluated TAG or TGA stop codon recoding across a dereplicated set of 9322 complete or nearly complete (≥90% complete) phage genomes from five different human and animal microbiome types. Specifically, we identified phages in gut microbiomes from people predicted to eat a westernized diet (samples: pregnancy cohort in Palo Alto CA, USA^23^, pregnancy cohort in Pittsburgh PA, USA^24^, residents of Norman OK, USA^25^, and residents of Bologna, Italy^26^), from people predicted to consume a non-westernized diet (samples: Hadza hunter-gatherers in Tanzania^27^, Matses hunter-gathers in Peru^25^, residents of Tunapuco, Peru^25^, arsenic-impacted adults in Eruani village, Bangledesh^18^, cholera patients and family members in Dhaka, Bangladesh^28^), and from baboons^29^, pigs^30,31^, cattle, and horses. We identified a set of candidate recoded phage genomes by looking for phage genomes that increased in coding density when genes were predicted with an alternative code instead of standard code (**Fig. 1A, Methods**). We then manually verified these putative alternatively coded (AC) genomes, arriving at a final set of 473 AC double-stranded DNA phage genomes. Previously stop codon recoding had only been found in phages with large genomes: crAssphages^17,20^ (95-190kb), jumbophages^19^(200-500 kb), and megaphages^18,21^ (>500kb to 660 kb). Here we identified complete AC phage genomes with the potential to circularize across a very wide diversity of sizes, ranging down to 14.7 kb (**Fig. 1B)**.

**Fig. 1:**
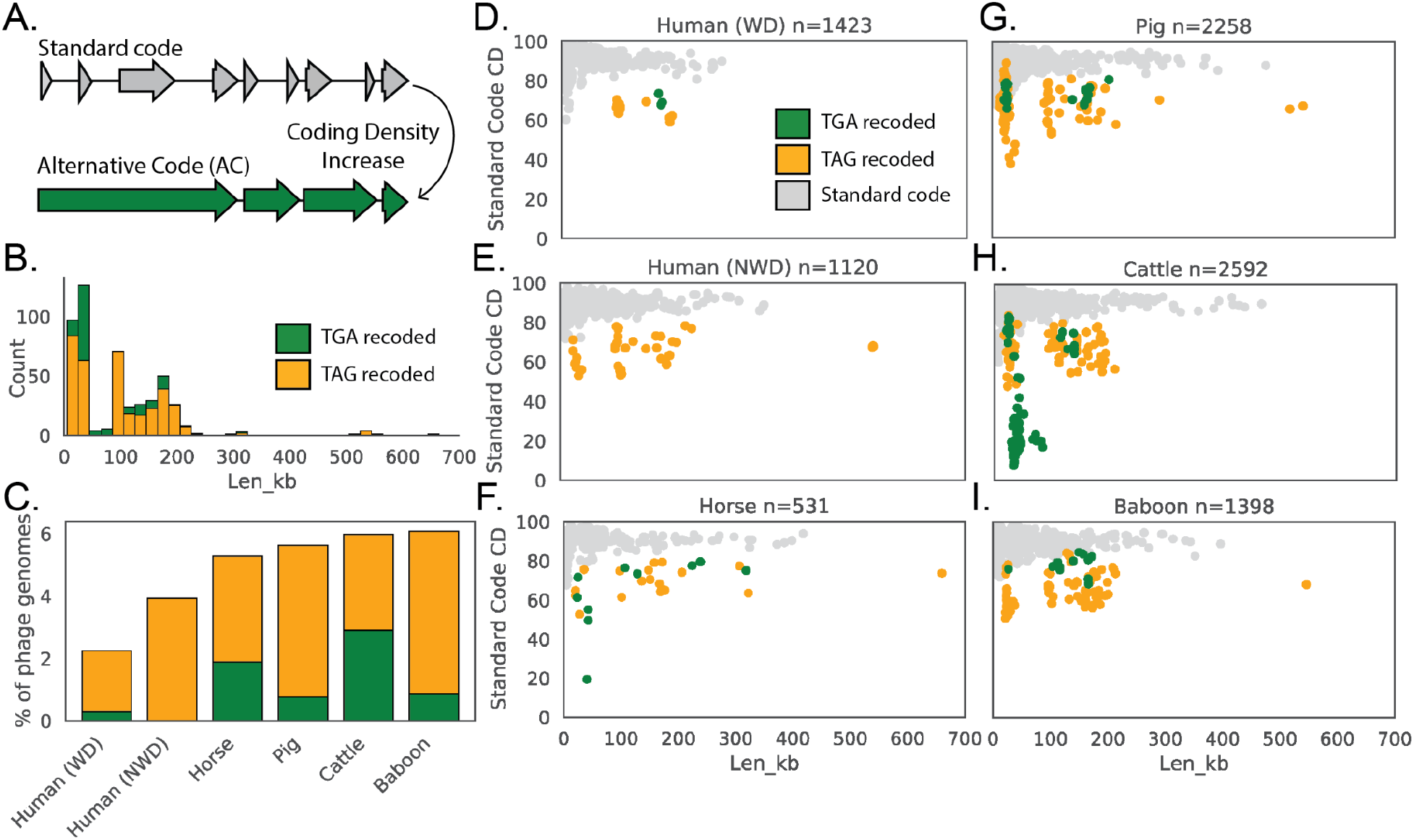
Recovery and identification of alternatively coded phages across human and animal microbiomes. **A**. A coding density increase between standard code and alternative code (AC) was used to identify putative AC phages, followed by manual confirmation of code. **B**. AC phage genomes spanned a wide size range from 14.7 kilobases (kb) to 660 kb. **C**. Abundance of AC phages varied from ∼2-6 % of the total phage population in the gut microbiome types surveyed in this study. WD = westernized diet, NWD = non-westernized diet. **D-I**. Phage genomes recovered from the indicated human or animal microbiome. The number of phage genomes (n) recovered after dereplication from each environment is indicated in the title of each plot. Individual phage genomes are represented by single points and plotted by size and coding density (CD) in standard code (code 11). In all plots, phage genomes have been dereplicated and are complete or near complete (>=90%). Symbol color represents genetic code (TGA recoding = green, TAG recoding = orange, standard code = grey). Len_kb = Length in kilobases.

We observed that TAG recoding is more common than TGA recoding (75% TAG recoded, 25% TGA recoded, **Fig. 1B,C**) in the tested metagenomes. While each gut microbiome type has AC phages present, alternative coding was observed at the lowest frequency in phage genomes recovered from the westernized human microbiome. Only 2.25% of phage genomes recovered from humans predicted to consume a westernized diet used an alternative code (**Fig. 1C)** and these AC phages span a narrow size range from 99-190 kb (**Fig. 1D**). In contrast, the other five microbiome types examined have higher frequencies of AC phages (up to 6.4% of genomes recovered from baboons), and these phages encompass a wide range of genome sizes (**Fig. 1C, E-I**). To test if AC phage genomes can be readily found outside of warm-blooded gut microbiomes, we also searched 2 reptilian (Giant Tortoise) gut metagenomes and found that 3% of the phage genomes recovered use recoded stop codons (**Fig. S1A)**. We conclude that alternative coding is a common feature of phage populations in the human and animal gut, and occurs in phages of diverse genome sizes.

### Diversity of alternatively coded phages

To understand the evolutionary relationships among families of AC phages, and also among AC phages and standard code phages, we constructed a phylogenetic tree of large terminase subunits from AC phages and their close standard code relatives (identified by vContact2^32^). We found many of these sequences form clades with high bootstrap support (≥95%)(**Fig. 2A**). To add the stop codon reassignment information onto this tree, we analyzed the alignments of terminase sequences with in-frame stop codons translated to X, and found that in most cases TGA aligned best with tryptophan (genetic code 4) and TAG aligned best with glutamine (genetic code 15). Interestingly, a subset of sequences use code 25, where TGA is reassigned to glycine. Some of these phages have Candidatus Absconditabacteria predicted as the host. Candidatus Absonditabacteria also uses code 25^15^, and we hypothesize that these code 25 phages are likely infecting code 25 hosts. In all other cases, the AC phage clades are predicted to infect common standard code gut phyla, Firmicutes and Bacteroidetes. The two previously characterized AC phage clades, crAssphage^20^ and Lak^18^ phage both infect Bacteroidetes hosts (*Bacteroides* and *Prevotella*, respectively).

**Fig. 2:**
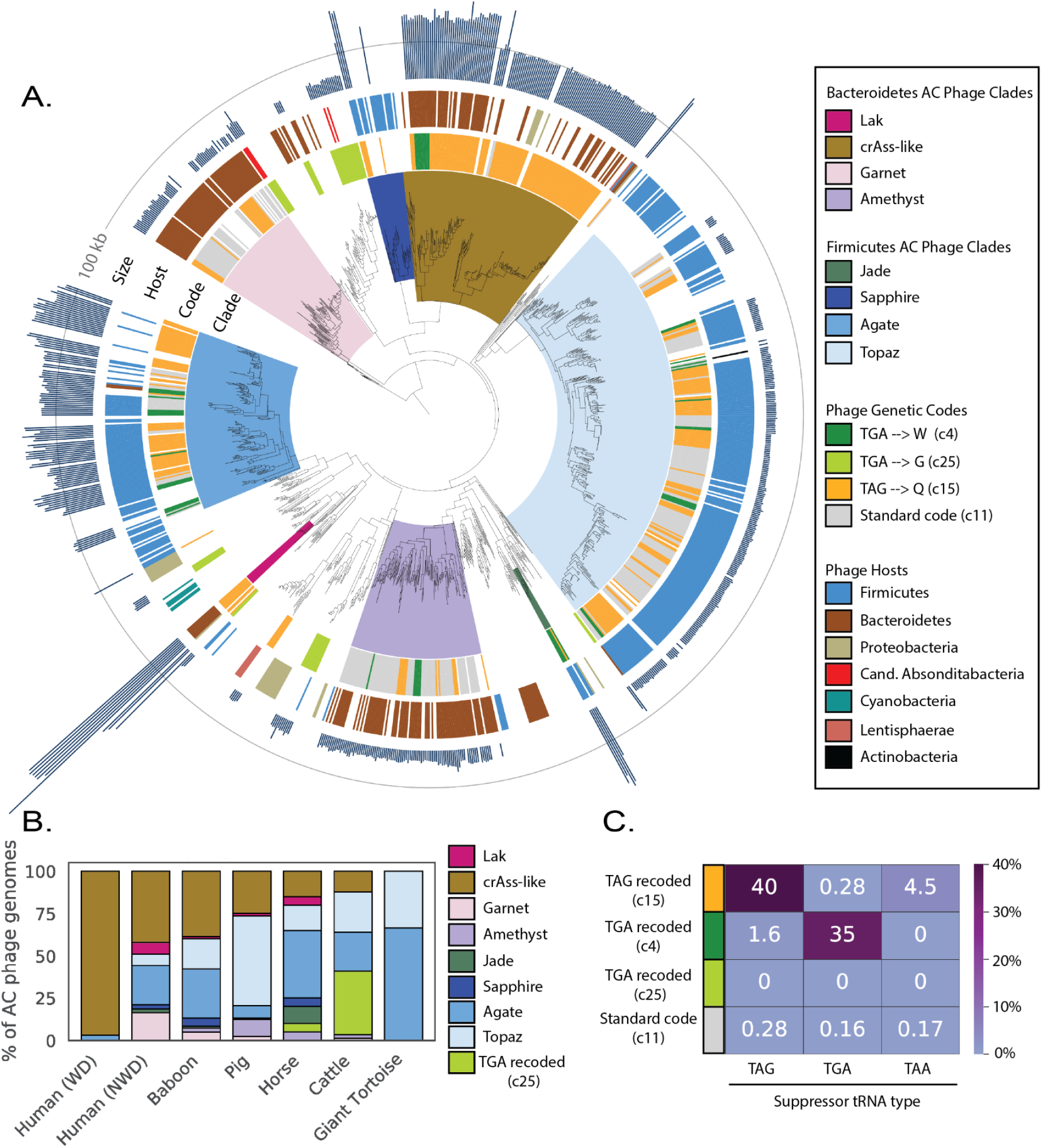
Phylogenetic reconstruction, environmental distribution, and suppressor tRNA use of phages with alternative genetic codes. **A**. The phylogeny of alternatively coded (AC) phages was reconstructed using large terminase sequences from a dereplicated set of complete or near-complete (>=90%) AC phages (n=444) and their close standard code relatives (n =258), as well as related proteins from Refseq r205 (n=410). Terminase sequences from eukaryotic herpesviruses (n=8) were used to root the tree. The inner to outer ring shows phage clade (>= 95% bootstrap support), genetic code for phages from this study, host phylum as predicted by taxonomic profiling and CRISPR spacer matches, and genome size with a grey line at 100 kilobases (kb) for scale. Genetic code and genome size were not included for Refseq proteins since some proteins were derived from prophages and/or incomplete phage genomes. **B**. Distribution of AC phages by clade across the 7 types of gut microbiomes evaluated in this study. WD = westernized diet, NWD = non-westernized diet. **C**. Heatmap of the percent of genomes of each genetic code that have tRNAs predicted to suppress translation termination at the TAG, TGA, or TAA stop codons.

Inspired by the historical designation of TAG and TGA as the amber and opal stop codons, we chose to name the six new clades of TAG and TGA recoded phages after other gemstones (Garnet, Amethyst, Jade, Sapphire, Agate, Topaz). Lak, crAss, Jade, Sapphire and Agate phages have larger than average mean genome length (562 ± 44kb, 210 ± 35 kb, 201 ± 22 kb, 154 ± 27 kb, ± SD) and Garnet, Amethyst, and Topaz phages have smaller than average genomes (34 ± 5kb, 34 ± 6kb, 22 ± 3 kb, ± SD). Genome size and predicted host phylum are relatively consistent within clades, however genetic code is not (**Fig. 2A, Table S1**). Of the eight AC phage clades identified here, four use code 11(standard code), code 15, and code 4 (**Table S1)**. Two clades use only code 15, and the other two use code 15 as well as either code 11 or code 4 (**Table S1**). These eight clades have uneven distributions across the environments analyzed here (**Fig. 2B**). Notably, the AC phage population associated with gut microbiomes of people from Europe and the US consists almost entirely of crAssphage, whereas the other gut microbiomes have higher diversity of AC phage types and higher abundance of Firmicutes-infecting AC phages (**Fig. 2B**).

Since most AC phages identified here are predicted to infect standard code hosts, they require some mechanism to change the meaning of a stop codon to a sense codon. Phage-encoded suppressor tRNAs, which decode stop codons, have been observed in AC phages before^17–21^. We predicted tRNAs in all phage genomes, and calculated the frequency at which genomes of each code (code 4, code 11, code 15, code 25) encoded suppressor tRNAs for the TAG, TGA, or TAA stop codons. We found a strong relationship between stop codon recoding and suppressor tRNA usage (**Fig. 2C**), although were only able to detect TAG suppressor tRNAs in 40% of TAG recoded genomes, and only 35% of TGA recoded (code 4) genomes had TGA suppressor tRNAs. It is likely that the fraction of phages that encode suppressor tRNAs is much higher than estimated here, as it would not be surprising if these phage tRNAs had noncanonical sequences or secondary structures that hindered their detection by existing methods. We also searched for phage-encoded release factors (RF), which terminate translation at stop codons and have been previously observed in Lak^18^ and crAss-like^17^ phages before. We identified RF2 (terminates translation at TAA and TGA) in six TAG recoded Lak phages and two TAG recoded Agate phages (**Table S2**). RF1, which terminates translation at TAG and TAA stop codons was identified in two TGA recoded Jade phages (**Table S2**). We also identified tryptophanyl tRNA synthetases in the same two Jade genomes (**Table S2)**, which we predict could ligate the amino acid tryptophan to the TGA suppressor tRNA, thus mediating the TGA → W code change. Our inability to identify release factors and tRNA synthetases mediating code change may be due in part to phage modification of bacterial tRNAs and translation machinery as well as the use of novel types of RNA molecules or enzymes by AC phages.

### Relationship to phages that use standard code

Our phylogenetic reconstruction of AC phages and their standard code relatives demonstrates that phages in the same clade may use different genetic codes. However, phages within the same clade may still be separated by significant evolutionary distances. To better understand the fine-scale relationships between phages that use different genetic codes, we calculated the average nucleotide identity (ANI) between all phages in our dataset to identify examples of closely related genomes that use different genetic codes. In a set of Agate genomes, we found phage genomes with greater than 80% ANI that use different genetic codes, and even found an example where a TGA recoded Agate phage (Cattle_ERR2019405_scaffold_1063) and a standard code Agate phage (pig_ID_3053_F60_scaffold_12) share greater than 90% ANI (**Fig. 3A,B**). This indicates that genetic code can change over remarkably short evolutionary timescales. A whole genome alignment between the TGA recoded Agate phage and its standard code relative shows accumulation of TGA codons in structural and lysis genes, but not the nearby DNA replication machinery (**Fig. 3B**).

**Fig. 3:**
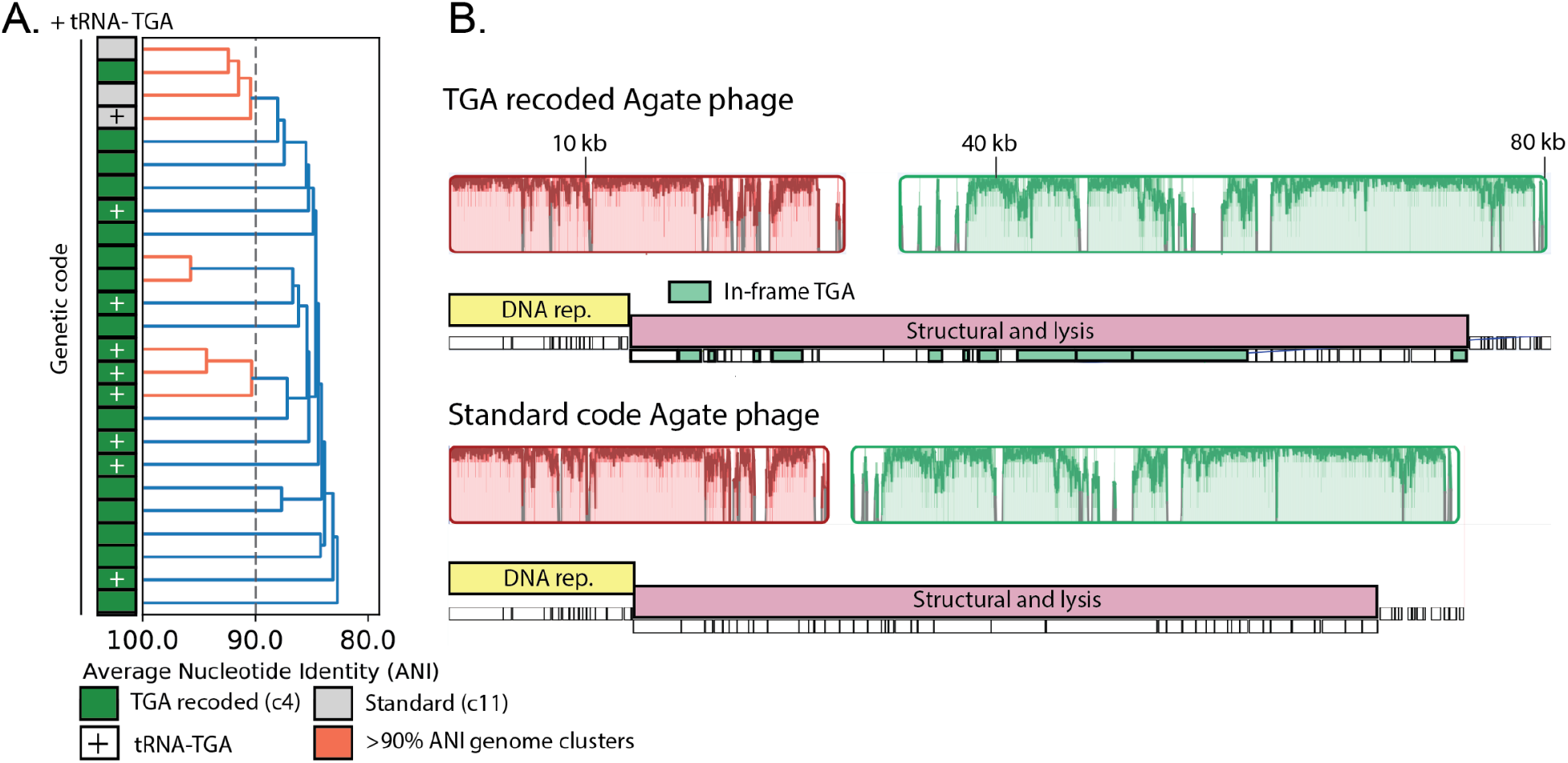
Close evolutionary relationships between standard code and alternatively coded phages. **A**. Dendrogram of average nucleotide identity (ANI) across a set of Agate phage genomes. Standard code (grey) and TGA recoded (green) phages share >80% ANI, and one cluster of >90% ANI genomes (orange) has both standard and TGA recoded genomes, indicating an extremely close evolutionary relationship. One standard code phage has acquired a TGA suppressor tRNA (+), potentially preceding code change. **B**. Global alignment of an 80 kilobase (kb) partial TGA recoded Agate genome (Cattle_ERR2019405_scaffold_1063) and a close standard code relative (pig_ID_3053_F60_scaffold_12). Homologous collinear sequences are shown with colored blocks (red and green here), where color corresponds to nucleotide alignment between the two genomes and lack of color represents lack of alignment. Genome structure for each phage is shown under the alignment graph, with DNA replication machinery represented as a yellow bar and structural and lysis genes with a pink bar. TGA stop codons have arisen in structural and lysis genes (individual recoded genes below in green) while DNA replication machinery has not accumulated stop codons.

This very close relative of TGA recoded Agate phages (pig_ID_3053_F60_scaffold_12, **Fig. 3B**) uses standard code and has only 4 out of 146 genes (2.7%) terminated with a TGA stop codon. In contrast, 34 genes use TAG and 108 genes use TAA. We hypothesized that divestment in TGA as a stop codon may have opened the door for its reemergence as a sense codon. To test for TAG or TGA stop codon loss at a wider scale, we surveyed stop codon usage across all standard code phages that are close relatives of AC phages (same relatives on phylogenetic tree, defined by vContact2^32^). These standard code close relatives strongly prefer the TAA stop codon over both the TAG and TGA stop codons (**Fig. S2A**, TAG vs. TAA: p = 8.03e-87, TGA vs. TAA: p = 2.97e-86, Mann-Whitney U Test). We also observed that TAG is rarer than TGA (**Fig. S2A**, p = 6.15e-11, Mann-Whitney U Test). This depletion of TAG and TGA is likely due to the significantly lower GC content (**Fig. S2B**, p = 4.08e-36, Mann-Whitney U Test) in these phages compared to standard code phages that are not close relatives of AC phages. Thus, stop codon loss driven by low GC content may be an evolutionary precursor to stop codon recoding in bacteriophages.

### Recoding as a potential regulator of cell lysis

Next, we explored the patterns of in-frame stop codon emergence across the genomes of alternatively coded phages. We observed that in code 25 phages predicted to infect code 25 hosts, an average of 90% of genes had in-frame TGA codons (**Fig. S3A**). However, in AC phage families predicted to infect standard code hosts, the mean fraction of genes using in frame stops ranges from 26% (Sapphire clade) to 71% (Garnet clade) (**Fig. S3A**). Less complete use of alternative code is consistent with a model where AC phages can translate early expressed genes using the standard code of their hosts (i.e., using standard bacterial machinery), and then produce molecules that enable the switch to translation of genes in which a stop codon is read as an amino acid.

We annotated in-frame stop codons across the genomes of representatives of each AC phage clade (**Fig. 4A-B, Fig. S4A-D, Fig. S5A-C**). Consistent with what has been previously observed, we saw that both the Lak^18^ and crAss-like^17,20^ genomes investigated here use alternative code for their “late” structural and lysis genes. Furthermore, we observed that Garnet, Amethyst, Jade, Sapphire, Agate, and Topaz phages also use alternative code for structural and lysis genes. In contrast, use of alternative code was variable in the DNA replication machinery. In crAss-like, Garnet, Amethyst, and Topaz phages, all the structural and lysis genes are encoded closely to one another as a single alternatively-coded genomic unit (**Fig. 4B, Fig. S4D, Fig. S4B-C**). In Jade, Sapphire, Agate, and Lak phages, the structural and lysis genes are in multiple alternatively-coded modules that are spread across the genome (**Fig. 4B, Fig. S4 A-C, Fig. S5A**). As structural and lysis proteins encoded with recoded stop codons cannot be expressed before the code change is manifested, stop codon recoding could effectively coordinate the timing of protein expression from related gene modules.

**Fig. 4:**
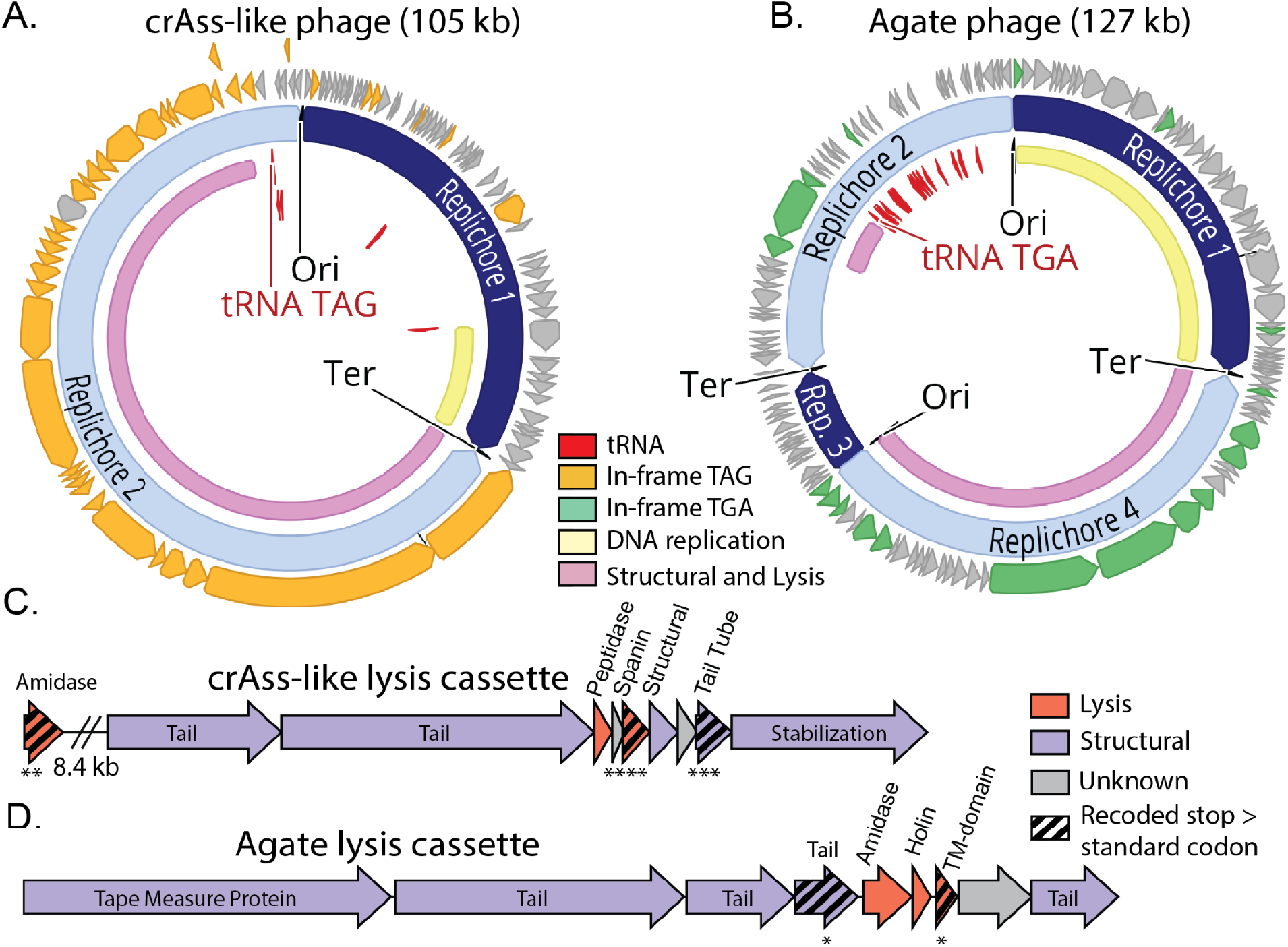
Preferential recoding of lysis-related genes in two distinct families of alternatively coded phages. **A-B**. Genomic maps of manually-curated representatives of crAss-like phages (js4906-23-2_S13_scaffold_20) and Agate phages (GiantTortoise_AD_1_scaffold_344_curated). TAG recoded genomes (**A**) contain genes with in-frame TAG codons (orange) while TGA recoded genomes (**B**) have genes with in-frame TGA codons (green). Suppressor tRNAs (red labels) are predicted to suppress translation termination at recoded stop codons. Regions of the genome encoding structural and lysis genes (pink) coincide with high use of alternative code. In these phages, DNA replication machinery (yellow) is encoded in standard code. Origins and termini were identified based on GC skew patterns indicative of bidirectional replication, and unique replichores are marked in alternating shades of blue. **C-D**. Genomic maps of highly-recoded lysis cassette neighborhoods from representative TAG-recoded crAss-like phages (**C**) and TGA-recoded Agate phages (**D**). Lysis genes (pink) as well as structural genes (purple) that were significantly biased towards use of in-frame stop codons (TAG in C, TGA in D) are marked with black striping. TM-domain = transmembrane domain. **** p ≤ 0.0001,*** p ≤ 0.001, ** p ≤ 0.01, * p ≤ 0.05, Benjamini-Hochberg p-value corrected Mann Whitney U Test.

We reasoned that by identifying the genes most biased towards use of recoded stop codons, we would identify the gene sets that were the most critical for phages to regulate in this manner. To this end, we searched for gene families that preferred to use the recoded stop codon to the standard code encodings of glutamine (code 15 phages, TAG → Q) or tryptophan (code 4 phages, TGA → W). We measured the codon preference in two distinct virulent phage types that were represented by a sufficiently large set of related genomes to enable gene-level statistics: ∼105 kb TAG-recoded crAss-like phages and ∼127 kb TGA-recoded Agate phages. While many genes in these phages have at least one in-frame stop codon (**Fig. 4A-B**), only a few gene families preferentially use the recoded stop codons at a statistically significant level (p < 0.05, Mann-Whitney U Test, corrected for multiple comparisons).

In the crAss-like phage genomes analyzed, only four gene families preferentially use TAG rather than CAG or CAA to encode Q (**Fig. 4C**). Two of these four families are essential components of the lysis cassette: a lysozyme type amidase (p = 6.82e-3) and a spanin, which is a critical regulator of lysis of gram-negative bacteria^33,34^ (p=9.00e-6). A tail tube gene family that is encoded two genes downstream (1.1 kb) of the spanin gene is also preferentially recoded (p = 7.59e-4). Having multiple strongly alternatively coded genes in the same transcript may amplify the stop codon mediated translation block. The fourth recoded gene family is of unknown function.

In the Agate genomes analyzed, three gene families preferentially use TGA instead of TGG to encode W (**Fig. 4D**). One of these is a group I intron endonuclease (p = 1.32e-3) that is inserted in an early gene. This self-splicing intron is expected to excise itself from the mRNA, but then in-frame stop codons should prevent homing endonuclease production until late in the infection cycle. A tail protein directly upstream of the lysis cassette (p = 4.18e-2) is also preferentially recoded, analogous to the tail tube gene in the crAss-like phages. A strongly recoded transmembrane domain protein (p = 4.64e-2) in the lysis cassette belongs to a family of transmembrane proteins that are assigned various lysis and lysis regulation related functions (holin, spanin, lysis regulatory protein, and ATP synthetase B chain precursor). We hypothesize a putative role for this protein in controlling lysis, potentially by depolarizing the cell membrane^35–38^.

Further supporting the link between lysis regulation and code change is the discovery of a “code change module” composed of a suppressor tRNA, a tRNA synthetase, and a release factor directly upstream of the lysis cassette in Jade phages (**Table S2, Fig. S6A-B**). These code-change related genes are all encoded in standard code, whereas the lysis genes directly downstream use alternative code (**Fig. S6A**). We anticipate that expression of these code change genes would drive expression of the lysis program. Overall, we propose that by changing the genetic code of the infected cell over time, these phages can use stop codon recoding to coordinate protein expression from related late genes and also to suppress misexpression of critical lytic gene products.

### Recoding in prophages as a potential lysogeny switch

So far we have described phage alternative genetic codes in the context of lytic growth. However, many phages are temperate, and integrate into their bacterial host chromosome as prophages. Excitingly, we found multiple examples of alternatively coded Garnet and Topaz prophages, two of which we analyzed in depth (**Fig. 5A-B**).

**Fig. 5:**
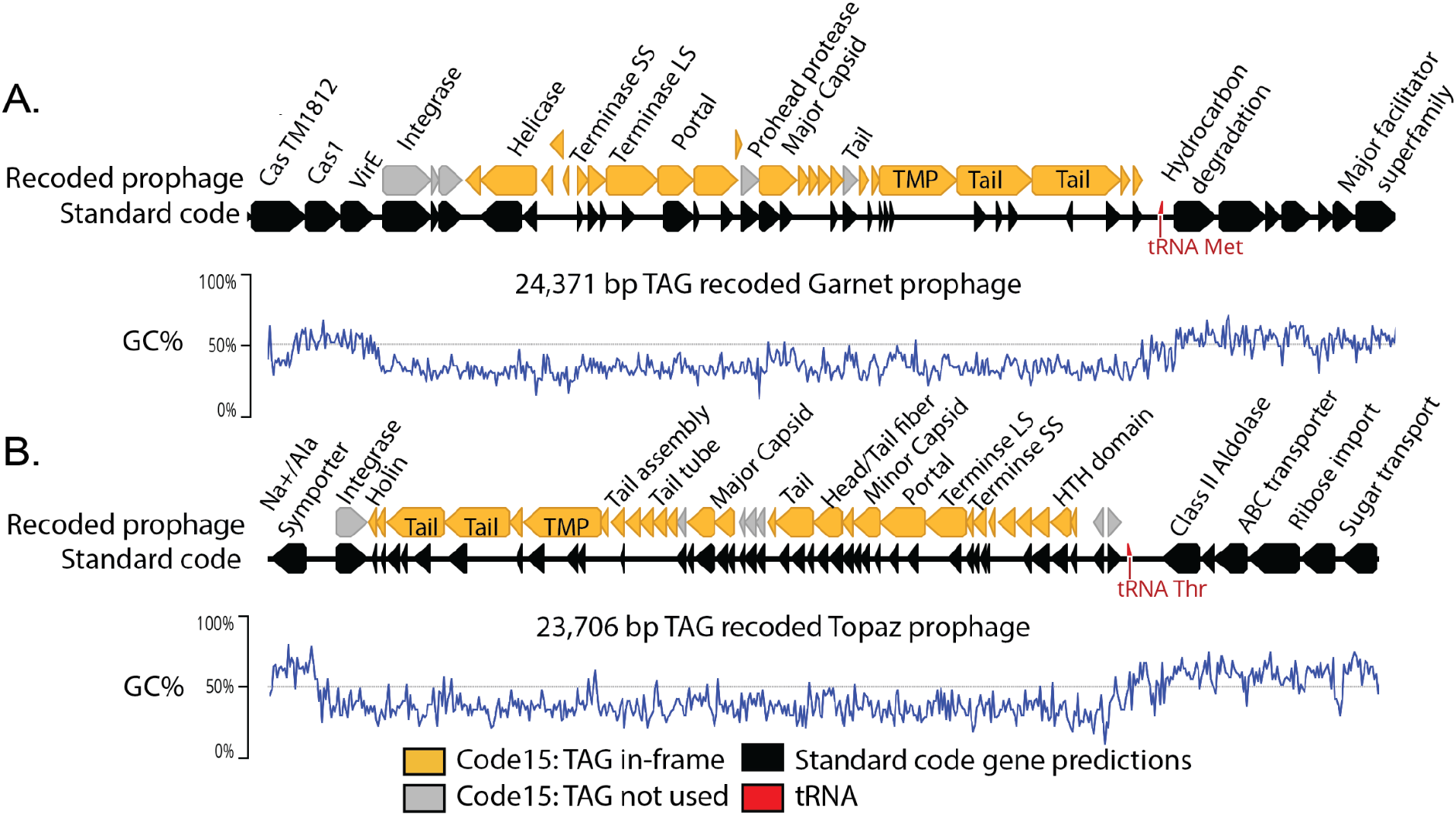
Alternatively coded prophages integrated into standard code bacterial genomes. **A**. Manually curated 24,371 bp TAG-recoded Garnet prophage integrated in a *Prevotella sp*. genome. **B**. A manually curated 23,706 bp TAG-recoded Topaz prophage integrated in a *Oscillospiraceae sp*. genome. In both A and B, the bacterial hosts use standard code (black gene predictions). Standard code results in highly fragmented gene predictions in the prophages, due to the high number of genes with in-frame TAGs (orange). In both A and B, the integrase is one of the few prophage genes that does not have in-frame TAG codons (grey). An increase in GC content (blue line) and transition from phage to bacterial gene content marks prophage boundaries. TMP = Tape Measure Protein.

The Garnet prophage is part of a 94 kb contig (SRR1747048_scaffold_47) belonging to a *Prevotella sp*. (Bacteroidetes) (**Fig. 5A)**. Notably, when we mapped reads to this prophage we observed that the sequencing read depth of the bacterial region was twice that of the integrated prophage (**Fig. S8A)**. As expected, a subset of the reads spanned the prophage, corresponding to *Prevotella* genomes that lack the integrated prophage. Thus, the exact prophage 24,371 bp genome could be defined.

The Topaz genome is part of a 36.9 kb contig (SRR1747065_scaffold_956) that is a close match to a contig from an *Oscillospiraceae sp*. (Firmicutes) (**Fig. 5B)**. In this case, sequencing reads coverage over the prophage region is ∼50 times higher than the flanking genome (**Fig. S7A)**. Many reads aligned with the ends of the prophage genome are discrepant at one end and the discrepant regions match the sequence at the other end of the prophage genome, indicating that these reads come from circularized sequences (**Fig. S7A)**. We infer the vast majority of phages in the sample were replicating and only a subset remained integrated at the time of sampling. Based on the sequence margins, we determined that the length of this prophage genome is 23,706 bp.

We also identified circular free phage genomes in our dataset that were nearly 100% identical to the Garnet and Topaz prophages. This supports our conclusion that these prophages represent actively-replicating genuine phages instead of defunct prophage genomes (**Fig. S9A-B**) and verifies the lengths determined from the read mapping analysis.

We noticed that while almost all of the prophage genes were extremely fragmented in standard code, the integrase genes did not contain in-frame stop codons (**Fig. 5A-B, Fig. S4D, Fig. S5B)**. Furthermore, we observed that all Garnet and Topaz integrase genes strongly avoided in-frame stop codons (Garnet: p=5.12e-3, Topaz: p = 1.229184e-16, Mann-Whitney U Test, corrected for multiple comparisons). We hypothesize that these phages are using stop codon recoding as a regulator of the lytic-lysogenic switch. In this scenario, the standard code translation environment of the host promotes expression of the integrase and establishment of lysogeny, with strong suppression of lytic genes. Likewise, a switch to alternative code would promote expression of lysis-related proteins during initial infection or prophage induction. Thus, genetic code may function as a mechanism to partition two distinct arms of the phage life cycle.

## Discussion

In this work, we show widespread use of recoded stop codons across eight distinct families of phages and prophages found in human and animal gut microbiomes. We hypothesize that the evolutionary progression from standard code to alternative code involves ancestral depletion of TAG and TGA stop codons, and propose a model where stop codon recoding is a post-transcriptional regulator of protein expression in phages and prophages (**Fig. 6**).

**Fig. 6:**
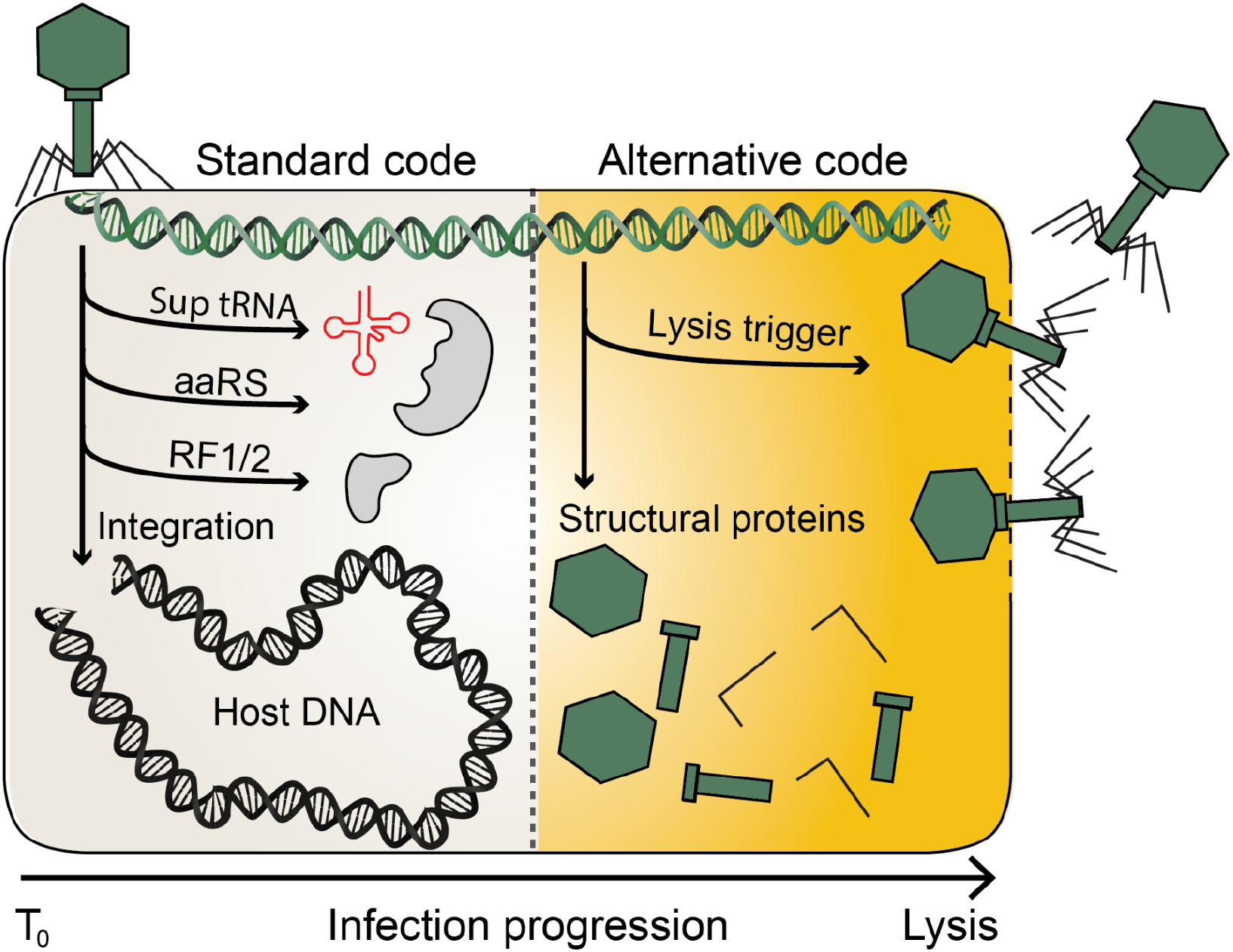
A model for of standard and alternative code use throughout the phage life cycle. Infection of a standard code host begins with the production of proteins from standard code compatible genes. In some phages, this is a route to integrase production and establishment of lysogeny. In other phages, this early phase involves the production of molecules involved in switching from standard to alternative code such as suppressor tRNAs (Sup tRNA), amino acyl tRNA synthetases (aaRS) and release factors (RF1/2). As the infection proceeds, alternatively-coded gene products initially suppressed by in-frame stop codons code can be produced. This allows for expression of phage structural proteins and ultimately triggers lysis.

A notable aspect of this study is that we identify stop codon recoding at a far higher rate than previously appreciated^39^. This is for two reasons: First, we surveyed a diverse set of microbiomes and so our analyses were not affected by the bias towards westernized human microbiomes that sometimes affects large scale microbiome analyses. Second, we restricted our analysis to complete and near-complete genomes, four of which were manually curated to ensure accuracy throughout. Since stop codon recoding is often only present in part of the genome, the recoded region of the genome may be greatly reduced or even entirely missing from an incomplete genome.

Using this large and diverse dataset of phages, we recovered extremely close relatives of AC phages, allowing us to describe what the immediate standard code relatives of AC phages look like. We propose an evolutionary route to recoding that begins with depletion of TAG or TGA stop codons in phages with low GC content. A rare stop codon is a useless but dangerous element: while not being used as a true stop, it can still cause damage as a nonsense mutation. Via acquisition of a means of code change such as a suppressor tRNA, in-frame stop codons can accumulate in positions that would have been previously lethal for the phage. Corroborating this model, we see TGA suppressor tRNA acquisition by standard code close relatives of TGA recoded phages (**Fig. 3A**). Also supporting this model is the observation that TAG stop codons are rarer than TGA stop codons in standard code relatives of AC phages, potentially explaining the higher prevalence of TAG recoding compared to TGA recoding. While we show a probable route from standard to alternative code, we acknowledge the possibility of evolutionary routes from alternative code back to standard code, although the mechanism for this transition remains unclear.

We propose that after in-frame stop codons are “detoxified” by the acquisition of suppressor mechanisms such as tRNAs, selection enriches or depletes stop codons across specific gene families to create patterns of codon use that can be harnessed as a form of gene regulation. We see diverse families of AC phages have all converged upon using recoded stop codons to encode lysis and structural proteins. This is consistent with more limited observations of structural gene recoding seen in Lak^18^ and crAsslike^17,20^ phages, and supports a model where the genetic code of the infected cell changes throughout the phage infection cycle. Dynamic codon use throughout the infection cycle has been demonstrated in T4-like phages that encode large tRNA arrays, where late-expressed genes have codon use aligned with the tRNA repertoire^40,41^. While the primary driver of tRNA use in T4 is replenishment of translational resources after host genome and transcriptome degradation^41^, this may also represent a mechanism to toggle translation efficiency of late genes throughout the phage life cycle. Stop codons inherently are codons with the lowest translation efficiency, and we hypothesize that use of recoded stop codons in late expressed genes represents an extreme version of codon based regulation in phages. We find two distinct lineages of virulent phages have both converged on preferential recoding of the lysis cassette, for which precisely timed expression is critical. Premature lysis aborts the phage life cycle and limits phage production, and some anti-phage immune systems even exploit this by forcing early lysis^38,42^. By encoding critical lysis regulators with in-frame stop codons, these phages block both accidental and host-forced premature expression of these proteins.

We also observe prophages with recoded stop codons integrated into standard code hosts, providing incontrovertible evidence that AC phages infect hosts with incompatible genetic codes. The long-term co-residence of two genomes with different genetic codes in the same cell has previously only been observed with alternatively coded organelles/endosymbionts and their associated nuclear genomes. We hypothesize that alternative coding may function as a lysis-lysogeny switch in these temperate AC phages. The decision to enter lytic growth or lysogeny is a critical point in the temperate phage life cycle, and phages have evolved elaborate regulatory mechanisms to precisely control this decision^43–45^. By encoding the lytic module and the lysogeny module in two different genetic codes, an AC phage would reduce the chance of gene products from one module interfering with the other. Recoding could also allow a phage to sense the presence of coinfecting or superinfecting phages that are using the same genetic code by responding to production of translation-related molecules such as suppressor tRNAs.

A final question is the cellular mechanisms to code change. While we identify suppressor tRNAs in 35-40% of AC phages, we rarely find phage-encoded tRNA synthetases that are predicted to recognize suppressor tRNAs and charge them with amino acids. We hypothesize that in some cases, the phages are relying on host tRNA synthetases to charge these suppressor tRNAs. We only observe TAG to glutamine and TGA to tryptophan reassignments in phages that use a code different from their host. This suggests that the bacterial glutaminyl and tryptophanyl tRNA synthetases are intrinsically susceptible to being repurposed by the phage to enact code change. Supporting this is literature demonstrating that TGA suppressor tRNAs are tryptophylated in *E. coli*^*46*^, and TAG suppressors likewise are glutaminylated^47^. A phage could potentially enact code change by stimulating this side reaction to become the dominant activity of the tRNA synthetase.

## Conclusion

Here, we surveyed phages for use of alternative genetic codes with reassigned stop codons. We find that stop codon recoding is widespread across diverse bacteriophages and prophages of the gut microbiome. Beyond the potential for stop codon recoding to play a previously unappreciated regulatory role in the phage life cycle, understanding alternative genetic codes in phages is crucial to our ability to detect and classify phage sequences. We predict that the cellular mechanisms of phage-induced genetic code change are rapid, efficient, and diverse across the phages of the natural world. Broadening our view of genetic code diversity in phages has the potential to greatly augment our understanding of basic phage biology and bacterial translation, as well as enhance synthetic biology strategies to design new genetic codes.

## Acknowledgments

We thank Yun Song, Jaime Cate, Kim Seed, Grayson Chadwick, Lin-Xing Chen, and Spencer Diamond for helpful discussions. We thank Ka Ki Lily Law and Jordan Hoff for technical support. This work was supported by a Miller Basic Research Fellowship to A.L.B, an NSF Graduate Research Fellowship to B.A-S (No. DGE 1752814), and NIH award RAI092531A, a Chan Zuckerberg Biohub award, and Innovative Genome Institute funding to J.F.B.

## Author Contributions

A.L.B and J.F.B. developed the project, led analyses, and wrote the manuscript with input from all authors. A.L.B, J.F.B, Y.C.L, R.S., and S.L. compiled the phage dataset. B.A-S assembled public metagenome data and provided support for phage genome analyses. Phage genomes were manually curated by J.F.B. A.L.J. and Y.C.L contributed to design of statistical analyses. J.M.S. contributed DNA samples from animal and arsenic-exposed human gut microbiomes.

## Data Availability

All supplementary materials, including genomes, proteins, and tRNAs of alternatively coded phages and relatives are available through Zenodo (https://doi.org/10.5281/zenodo.5275335).

## Methods

### Phage prediction

Phage prediction tools Seeker^48^ (predict-metagenome) and VIBRANT^49^ were run on assembled metagenomes (contigs > 5kb) using default settings. CheckV^50^ (end-to-end) was run on predicted phages and trimmed proviruses to evaluate completeness and quality. Contigs evaluated as low quality by both CheckV and VIBRANT were removed from analysis. Contigs < 100 kb with viral genes > host genes and contigs > 100 kb with < 20% host genes were maintained as high confidence phages.

### Phage dereplication

Phage scaffolds for each ecosystem were dereplicated at 99% ANI using the dRep^51^ dereplicate module (-sa 0.99 --ignoreGenomeQuality -l 5000 -nc 0.5 --clusterAlg single -N50W 0 -sizeW 1).

### Identification of phage genomes with recoded stop codons

Prodigal^52^ (single mode) was used to predict genes on dereplicated >= 90% complete phage genomes using genetic codes 4, 11, and 15. Coding density was calculated by summing the length of genes for each contig and dividing by the total contig length. Contigs 5-100 kb that had an increase of greater than 10% coding density in code 4 or code 15 relative to code 11 were tentatively assigned that genetic code, as were contigs >= 100 kb with a coding density increase >5%. All code assignments were confirmed by manual analysis of each contig. If the alternative genetic code resulted in more contiguous operon structure, reduced strand switching, correct-length genes (as checked by blastp^53^ against NCBI database), and did not result in gene fusions (as checked by blastp against NCBI database) the phage was confirmed as alternatively coded.

### Structural and functional annotations

Coding sequences predicted by prodigal using genetic code 4 for TGA recoded phages and code 15 for TAG recoded phages. HMMER^54^ (hmmsearch) was used to annotate the resulting sequences with the PFAM, pVOG, VOG, and TIGRFAM HMM libraries. tRNAs were predicted using tRNAscan-s.e. V.2.0 in bacterial mode^55^.

### Host prediction

A combination of CRISPR spacer analysis and taxonomic classification were used to predict putative host phyla. Contigs with a minimum length of 5 kb from the human and animal metagenomes analyzed in this study were searched for CRISPR spacers using minCED^56^. blastn short was used to identify matches between phage and spacer of >90% identity and >90% spacer coverage. Taxonomic profiling was performed by using DIAMOND^57^ (fast mode, e = 0.0001) to search all phage proteins against a custom version of the UNIREF100 database that retained NCBI taxonomic identifiers. tRep^58^ was then used to profile the taxonomy of each phage contig. For each contig, the bacterial phylum with most hits was considered to be the putative host, but only if that phylum had more than 3x hits than the second most common phylum^19^. In almost every case, the CRISPR spacer analysis and the taxonomic profiling agreed on the phage host phyla. In the rare cases that these analyses were not in agreement, the host phyla was considered unknown.

### Phage genome clustering by average nucleotide identity (ANI)

Our total dataset of 9422 non-dereplicated phage scaffolds from all ecosystems was augmented with 1428 phage genomes from other animal/human microbiomes from ggkbase, and the genomes clustered using dRep^51^ compare module (-sa 0.8 -pa 0.8 -nc .1 --clusterAlg single). Whole genome alignment was visualized using Mauve^59^ (progressiveMauve algorithm).

### Phage clustering with Vcontact2

Phages scaffold from the dereplicated dataset of >= 90% complete phage scaffolds for each ecosystem were clustered into viral clusters with Refseq viruses using Vcontact2^32^ (--rel-mode ‘Diamond’ --db ProkaryoticViralRefSeq201-Merged --pcs-mode MCL --vcs-mode ClusterONE). Standard code phages that were in the same viral cluster (VC) as at least one alternatively coded phage were considered to be close relatives of alternatively coded phages.

### Phylogenetic analysis of large terminase subunit of alternatively coded phages and standard code relatives

Terminases were found using two rounds of HMM-based classification. Proteins were initially annotated using PFAM, pVOG, VOG, and TIGRfam HMMs. This did not result in complete recovery of terminases for all phages of interest. To increase sensitivity, we clustered proteins into subfamilies using MMseqs^60^ (-s 7.5, -c 0.5, -e 0.001), and used HHblits^61^ to generate hmms of each subfamily based on alignments generated with the MMseqs result2msa parameter. We used HHSearch ^62^ (-p 50 -E 0.001) to perform an HMM-HMM comparison with the PFAM database. We then identified subfamilies with a best hit to large terminase HMMs with a >95% probability. Putative terminase subfamilies with a low number of primary terminase annotations were confirmed by blastp against the NCBI database. If subfamily members had hits to terminases in known phages, we considered the subfamily to be a true terminase subfamily. In rare cases, the terminase gene was fragmented due to assembly error or mobile intron insertion. In these cases we chose the larger of the gene fragments for downstream analysis. Terminases from AC phage and these standard code relatives (from vContact2^32^) were searched against the Refseq protein database using blastp, retaining the top 10 hits per protein. The recovered Refseq proteins were dereplicated at 90% using CD-HIT^63^. AC phage, standard code relative, and deprelicated Refseq terminases were combined and aligned using MAFFT^64^, and the alignment trimmed with trimAL^65^ (-gt 0.5). IQ-TREE^66^ was used to build a tree using the VT+F+R10 model and ultrafast bootstrap with 1000 iterations. Tree was visualized using iTol^67^.

### Codon preference analysis

#### crAss and Agate analysis

ANI-based genome clustering showed high representation of a lineage of TGA recoded ∼127 kb Agate phages as well as a lineage of TAG recoded ∼105 kb crAssphages, which were chosen for further analysis. For each phage lineage, genes were clustered into families created using a two step protein clustering method. First, proteins were clustered into subfamilies using MMseqs^60^ (-s 7.5, -c 0.5, -e 0.001), and HHBlits^61^ was used to generate HMMs of each subfamily based on alignments generated with the MMseqs result2msa parameter. Thes HMMs were then compared to one another using HHBlits (-p 50 -E 0.001). MCLclustering (--coverage 0.70 -I 2.0 --probs 0.95) was used to generate families from the HMM-HMM comparisons. Wilcoxon rank sum test was used to evaluate protein families that preferred the in-frame stop codon to the standard coding for the recoded amino acid. The Benjamini-Hochberg p-value correction was used to correct for multiple hypothesis testing. For TGA → W recoded phages, TGA occurrence was compared to the occurrence of the standard codon for Trytophan (TGG). For TAG → Q recoded phages, TAG occurrence was compared to the occurrence of the standard codons for Glutamine (CAG, CAA). Proteins were annotated by PFAM, pVOG, VOG, and TIGRFAM as well as BLAST searches against the NCBI database. In some cases, the HHPred webserver^68^ and the Phyre2 webserver^69^ were used to augment initial annotations. Gene neighborhoods were visualized using Clinker^70^.

#### Garnet and Topaz analysis

Garnet and Topaz proteins were clustered into families using the two step method detailed above. We identified the integrase families for each phage clade using PFAM, pVOG, VOG, and TIGRfam HMM annotations. We observed that the majority of the integrase genes had zero in-frame stop codons. A few genes had one in-frame stop, and when we examined alignments of the integrase families we found that in all cases the in-frame stop was in a N or C terminal extension of the protein. We believe that this corresponds to incorrect start codon prediction (N terminal extensions) or legitimate use of the codon to terminate the integrase gene (C terminal extensions). To test if this depletion of in-frame stop codons in the integrase genes occurred at a rate higher than would be expected to occur by chance, we used Wilcoxon rank sum test to evaluate all protein families in each phage clade for avoidance of in-frame stop codons relative to standard codons for Glutamine (CAG, CAA) or Tryptophan (TGG). The Benjamini-Hochberg p-value correction was used to correct for multiple hypothesis testing. We found that for both Garnet and Topaz phages, the integrase gene families avoided in-frame stop codons at a rate higher than expected by chance alone (Garnet: p=5.12e-3, Topaz: p = 1.229184e-16).

#### Origin and terminus determination via GC Skew

GC skew (G-C/G+C) and cumulative GC skew were calculated across the phage genome^71^. This allowed us to predict origins of replication, replication termini, and define individual replichores. We observed a variety of replication styles: double origin bi-directional replication, single origin bi-directional replication, and unidirectional replication. We also observed GC skew patterns of unknown significance.

## Main Figure Legends

**Fig. S1:**
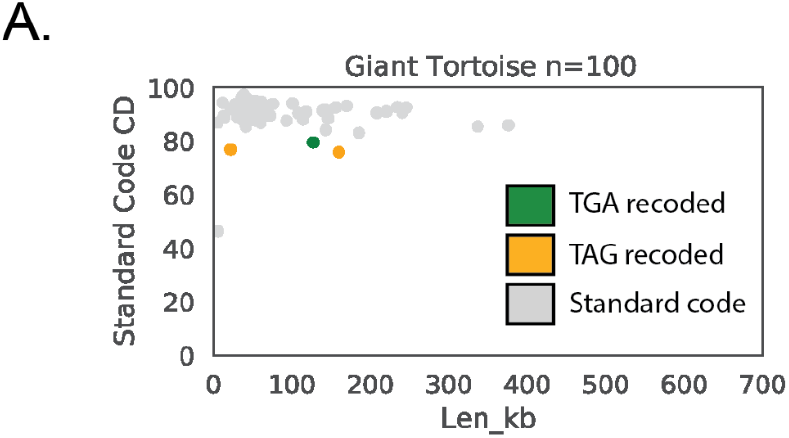
Identification of alternatively coded phages in the Giant Tortoise microbiome. **A**. Dereplicated complete or near complete (>=90%) phage genomes from the Giant Tortoise gut microbiome. Phages are plotted by size and coding density (CD) in standard code (Code11). A signature of stop codon recoding is low coding density in standard code. Phages that have recoded the TGA stop codon are indicated in green, and phages that have recoded the TAG stop codon are indicated in orange.

**Fig. S2:**
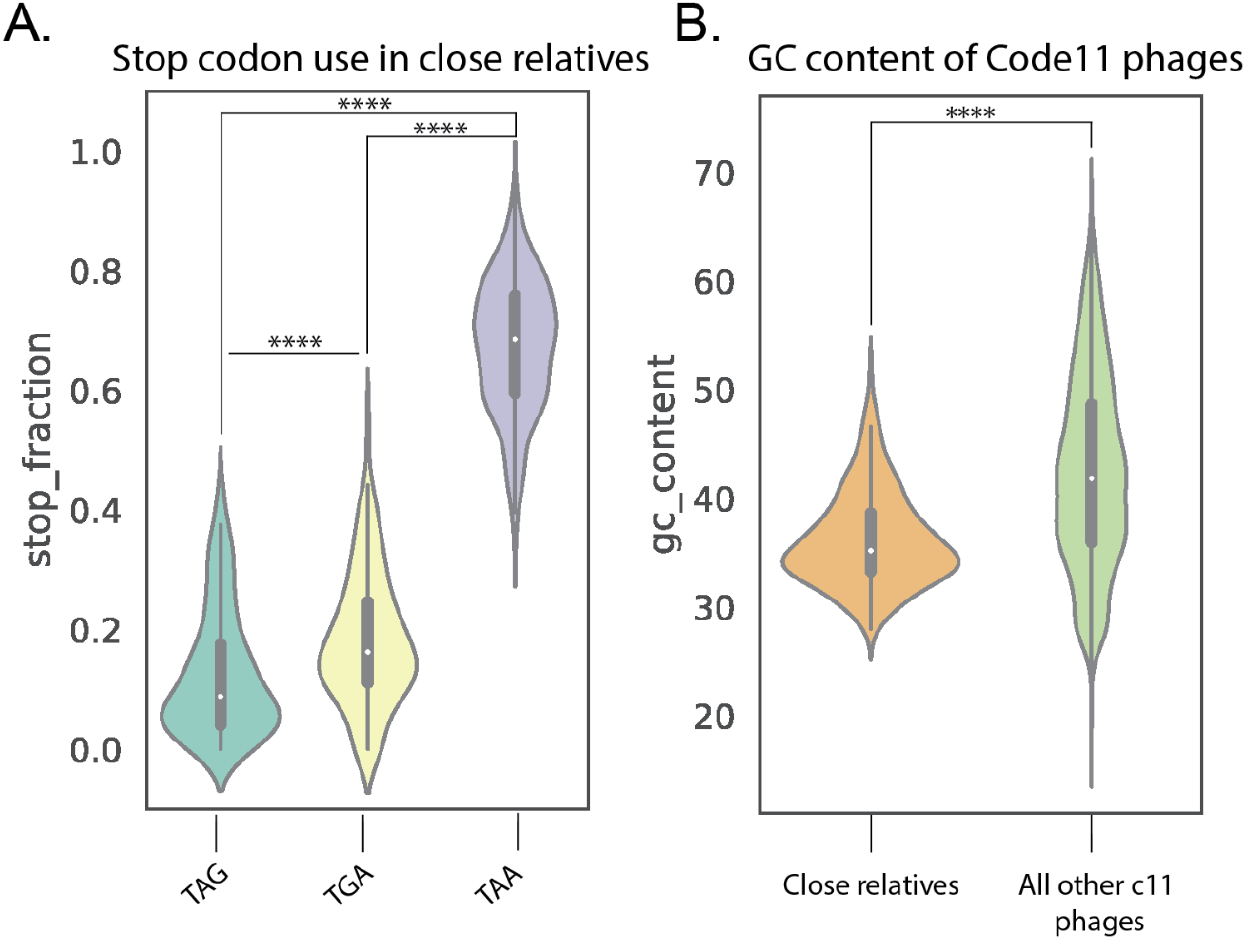
Stop codon use and GC content of standard code phages that are close relatives of phages that use alternative genetic codes. **A**. Close relatives of alternatively coded (AC) phages (n = 261) strongly prefer the TAA stop codon over both the TAG and TGA stop codons (TAG vs. TAA: p = 8.03e-87, TGA vs. TAA: p = 2.97e-86. The TAG stop codon was also depleted relative to TGA (p = 6.15e-11) in these phages. **B**. Close relatives of alternatively coded phages have a lower mean GC content relative to all other standard code phages (n = 8687, p = 4.08e-36). **** p ≤ 0.0001, Mann-Whitney U test. Close standard code relatives were identified using vContact2.

**Fig. S3:**
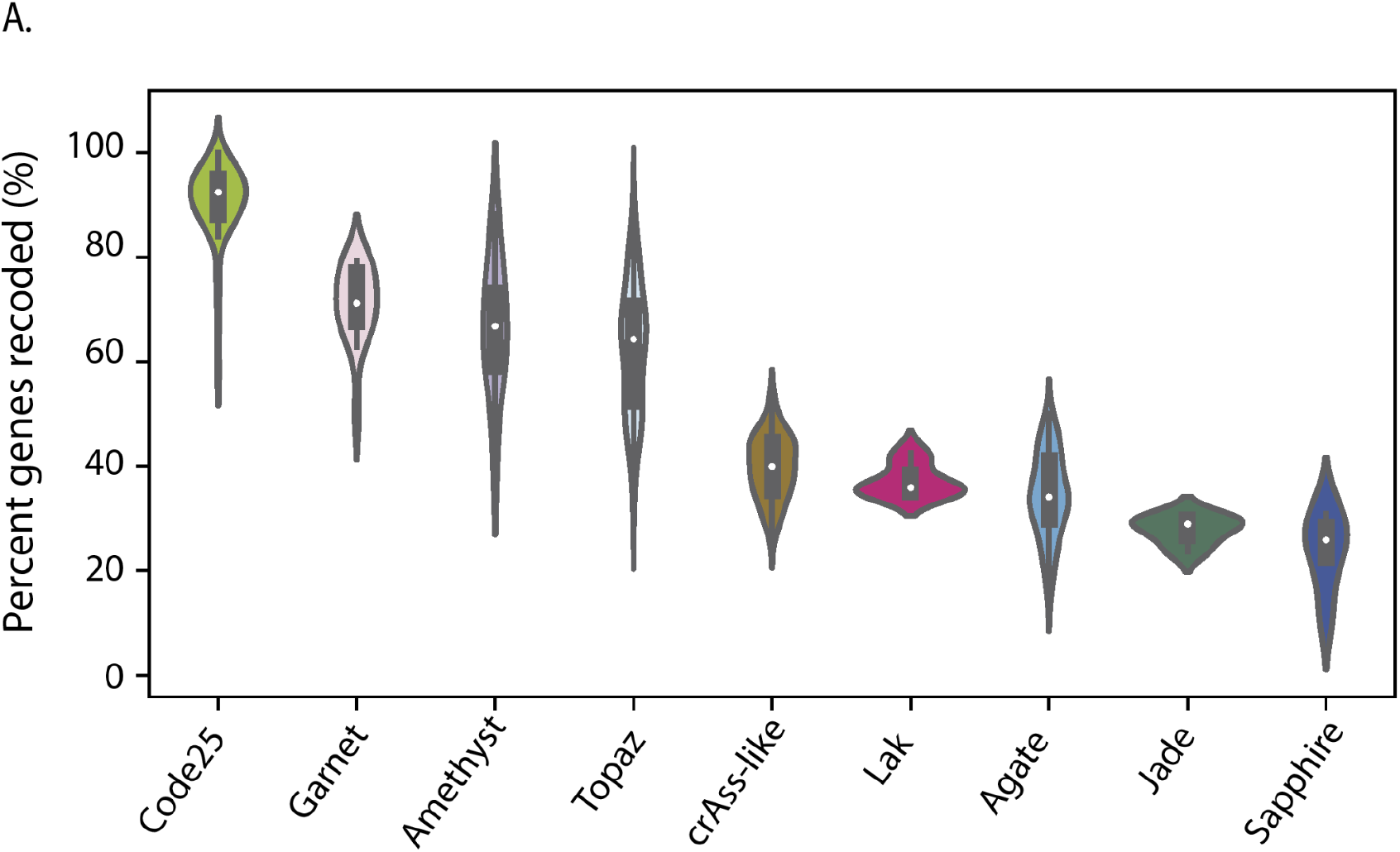
Percent of total genes that have recoded stop codons across different clades of alternatively coded phage. **A**. If a gene has at least 1 in-frame stop codon, it is considered recoded. Code 25 phages have the highest percentage of recoded genes, consistent with the hypothesis that they are infecting code 25 bacteria. In this scenario, these code 25 phages have fully adapted their genetic code to their host. The other phage types have lower percentages of recoded genes, consistent with the hypothesis that they are infecting standard code bacteria. We predict these phages switch from using standard code to alternative code part-way through their infection cycle.

**Fig. S4.**
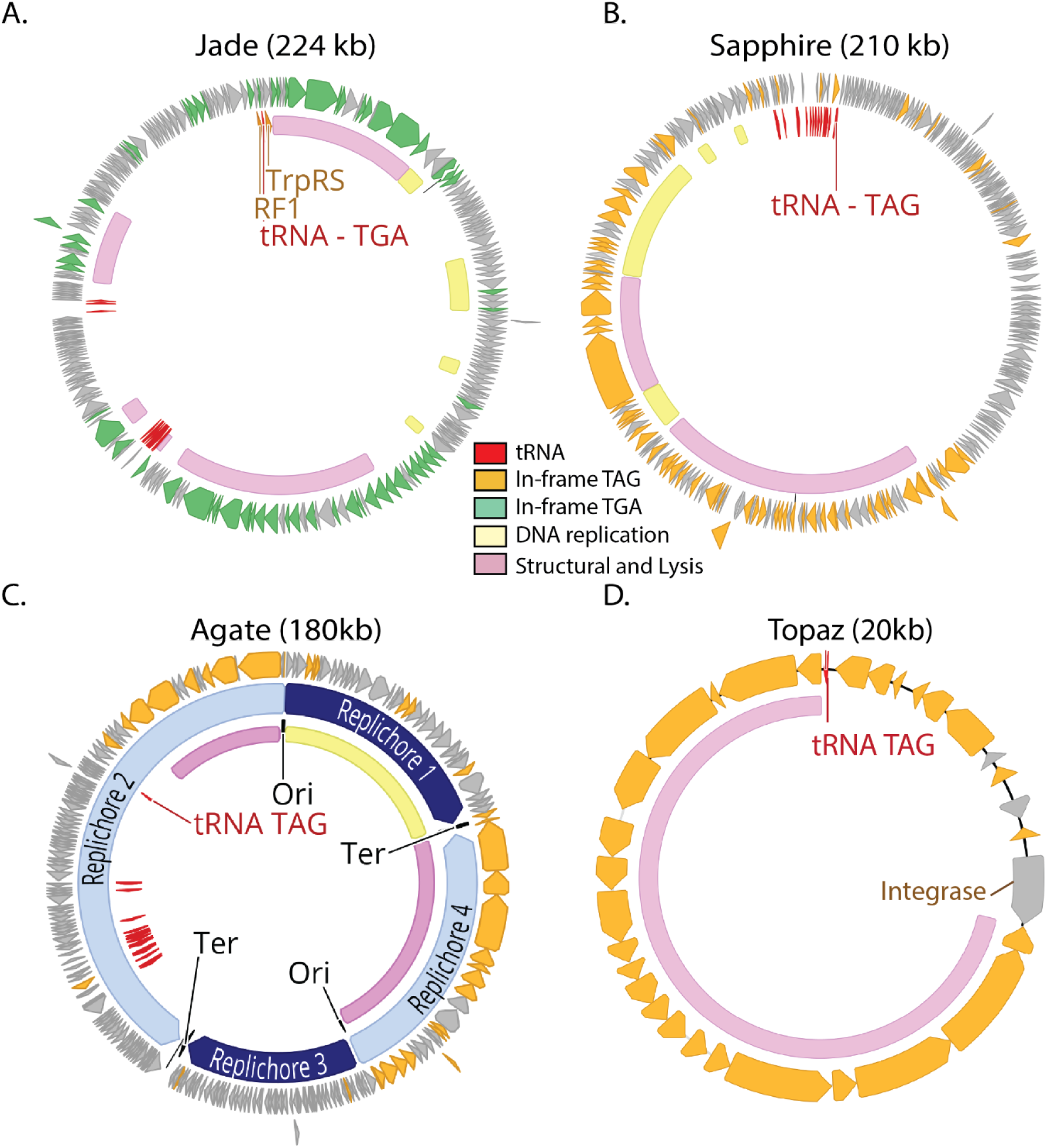
Genomic maps of representatives of Jade, Sapphire, Agate and Topaz clades. **A-D**. TGA recoded genomes (A) contain genes with in-frame TGA codons (green) while TAG recoded genomes (B-D) have genes with in-frame TAG codons (orange). Suppressor tRNAs (tRNA TGA or tRNA TAG, red) are predicted to suppress translation termination at TGA and TAG stop codons, respectively. In all cases, regions of the genome encoding structural and lysis genes (pink) coincide with high use of alternative code. Contrastingly, genes involved in DNA replication (yellow) are variably encoded in alternative code. Genomes with a GC skew patterns indicative of bidirectional replication and clear origins and termini (Agate, C) have unique replichores marked in alternating shades of blue. Genomes with unidirectional replication (Topaz, D) or unclear GC skew patterns (Jade, Sapphire, A-B) have no replication-related annotation. In some cases, unique or interesting genes have been noted with text. Clade representatives: Jade = JS_HF2_S141_scaffold_159238, Sapphire= SRR1747018_scaffold_13, Agate = Cattle_ERR2019359_scaffold_1067472, Topaz = pig_ID_1851_F40_2_B1_scaffold_1589.

**Fig. S5.**
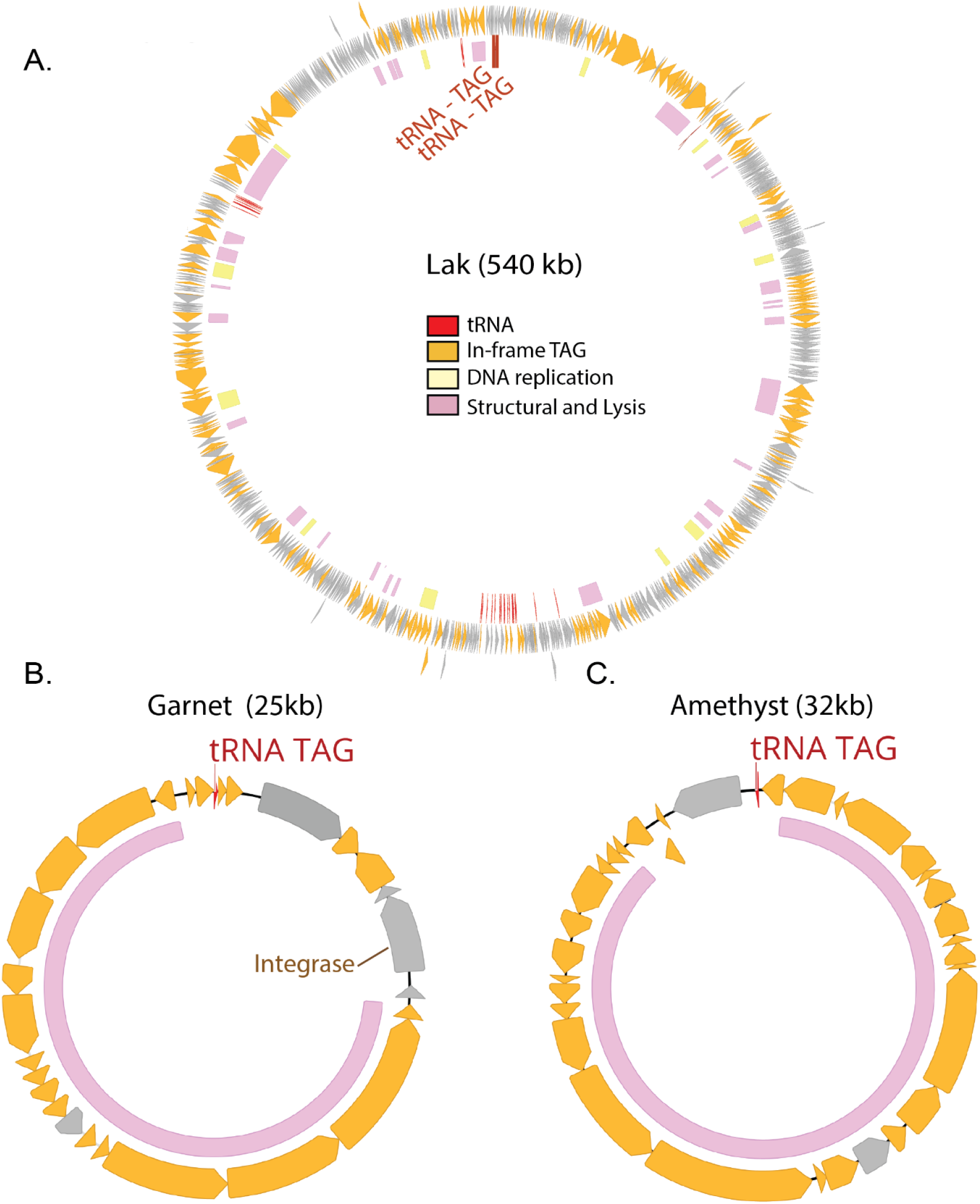
Genomic maps of representatives of Lak, Garnet, and Amethyst clades. **A-C**. TAG recoded genomes have genes with in-frame TAG codons (orange). Suppressor tRNAs (tRNA TAG, red) are predicted to suppress translation termination at TAG stop codons. Regions of the genome encoding structural and lysis genes (pink) coincide with high use of alternative code. In Lak phage (A), genes involved in DNA replication (yellow) are mostly encoded in alternative code. Origins and termini are unmarked in these genomes as we were unable to define clear replichores for Lak (A) and Garnet and Amethyst (B-C) appear to utilize unidirectional genome replication based on GC skew patterns. In some cases, unique or interesting genes have been noted with text. Clade representatives: Lak = C1--CH_A02_001D1_final, Garnet = pig_ID_3640_F65_scaffold_1252, Amethyst = pig_EL5596_F5_scaffold_275).

**Fig S6.**
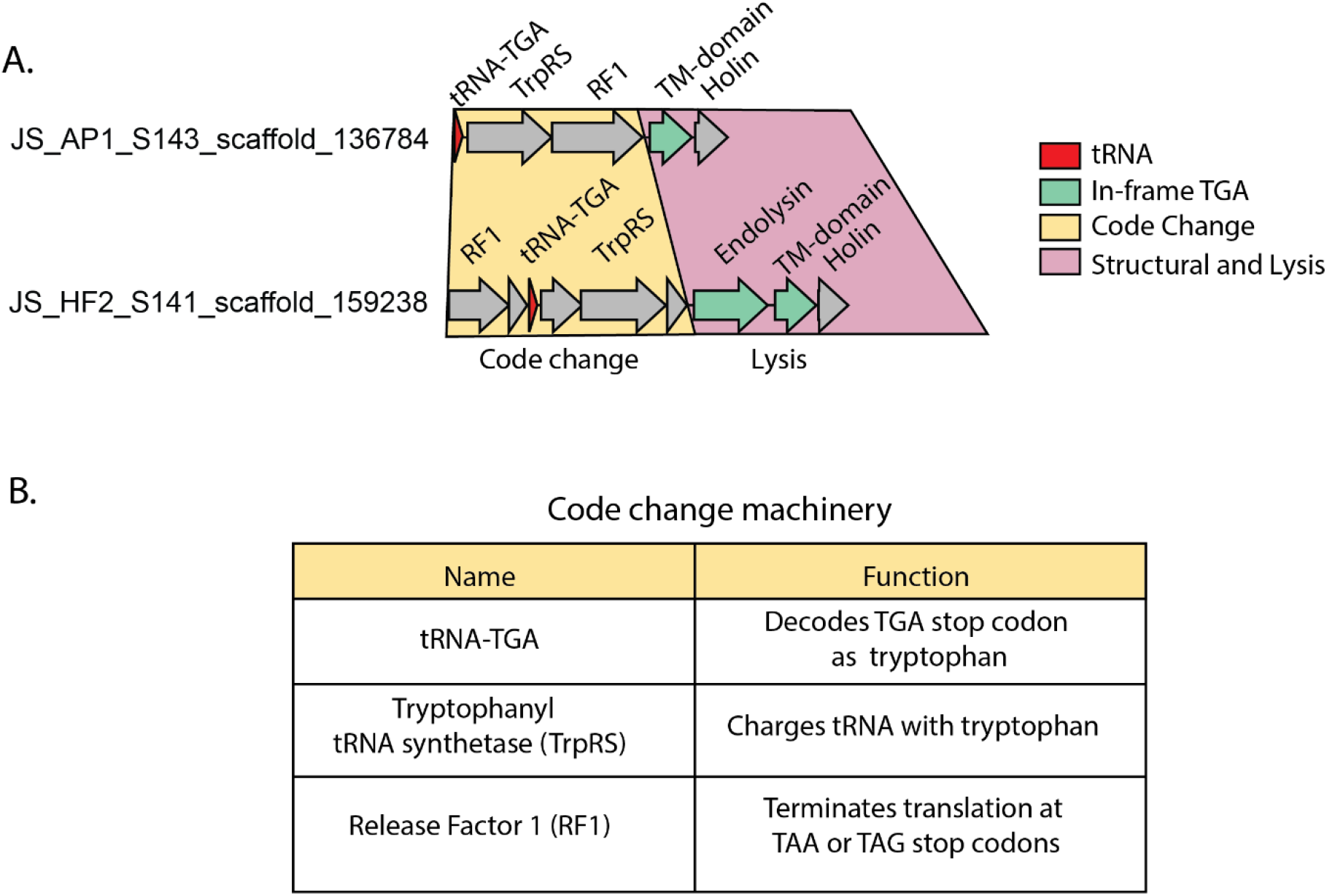
Code change machinery in two TGA-recoded Jade phages. **A**. An operon implicated in changing the genetic code from standard code (TGA = Stop) to code 4 (TGA = W) is directly upstream of the lysis cassette. The code change operon itself is encoded in standard code, while some genes in the lysis cassette have in frame TGA codons (green). TrpRS = Tryptophanyl tRNA synthetase, RF1 = Release Factor 1, TM-domain = Transmembrane domain. **B**. Functions of genes implicated in changing the genetic code of the infected Firmicutes cells. The tryptophanyl tRNA synthetase is predicted to charge the TGA suppressor tRNA with tryptophan. Release Factor 1 expression is predicted to stimulate translation termination at TAG and TAA while the TGA tRNA suppresses termination at TGAs.

**Fig. S7.**
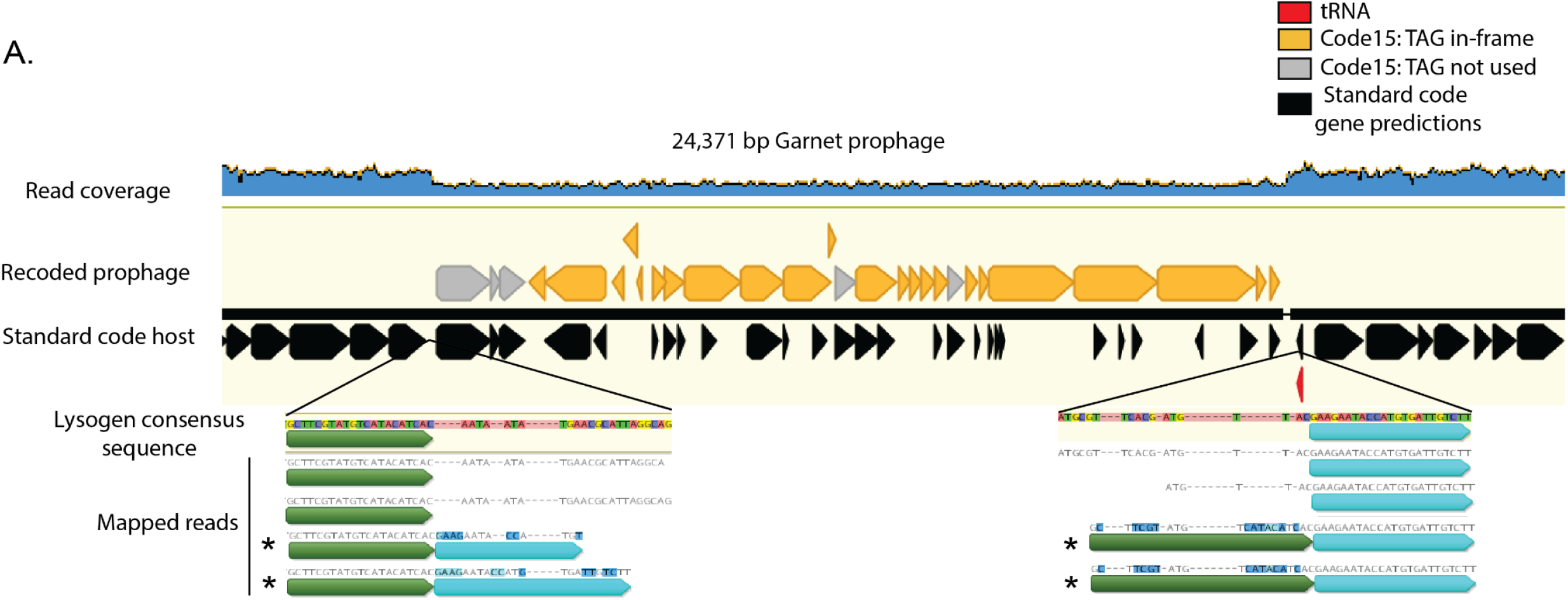
Read mapping to Garnet lysogen. **A**. Reads were mapped against a manually curated Garnet lysogen. Read coverage for the Prevotella sp. DNA is ∼2x higher than the read coverage of the Garnet prophage, indicating that the bacterial population in this sample is incompletely lysogenized. Supporting this conclusion are paired reads that span the length of the prophage (not shown), as well as some individual reads which show imperfect mapping to the lysogen consensus sequence (marked with asterisk). These reads represent the contiguous bacterial sequence. Identical sequence blocks are indicated with color.

**Fig. S8.**
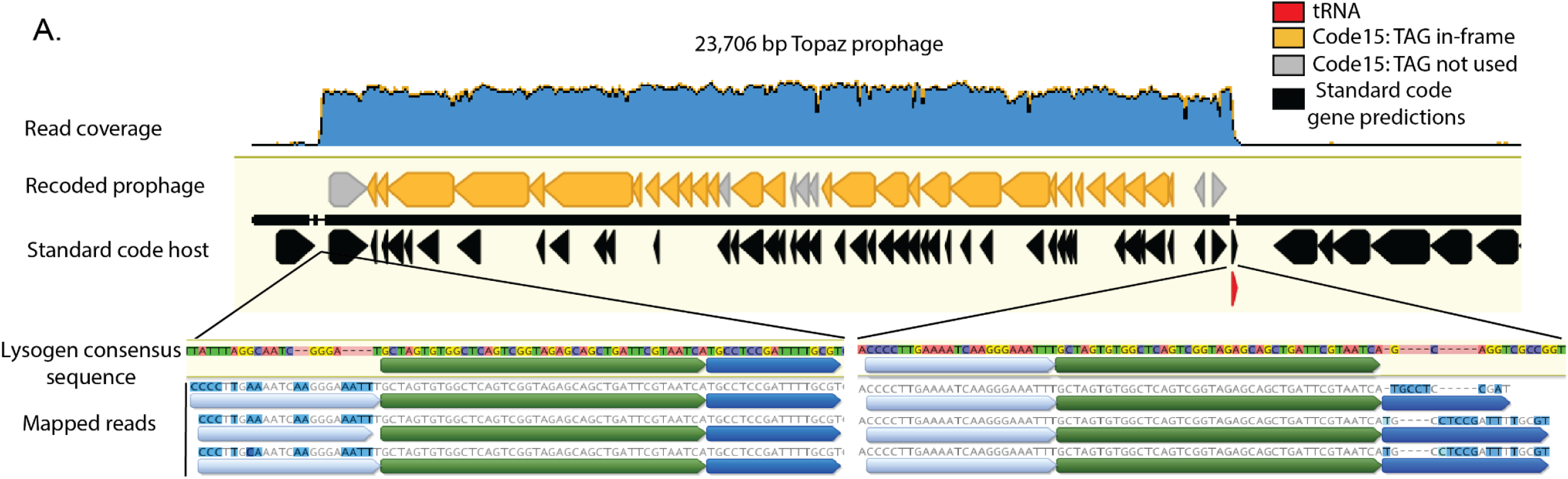
Read mapping to Topaz lysogen. **A**. Reads were mapped against a manually curated Topaz lysogen. Read coverage for the integrated Topaz phage genome is ∼50x higher than the neighboring *Oscillospiraceae sp*. sequence. This indicates that the phage is actively replicating in this sample. Supporting this conclusion are paired reads that span the length of the prophage (not shown), as well as individual reads which show imperfect mapping to the lysogen consensus sequence and contain sequences identical to the other side of the phage genome (light blue and dark blue sequence blocks). These reads are derived from circularized phage genomes in the sample. Identical sequence blocks are indicated with color.

**Fig. S9.**
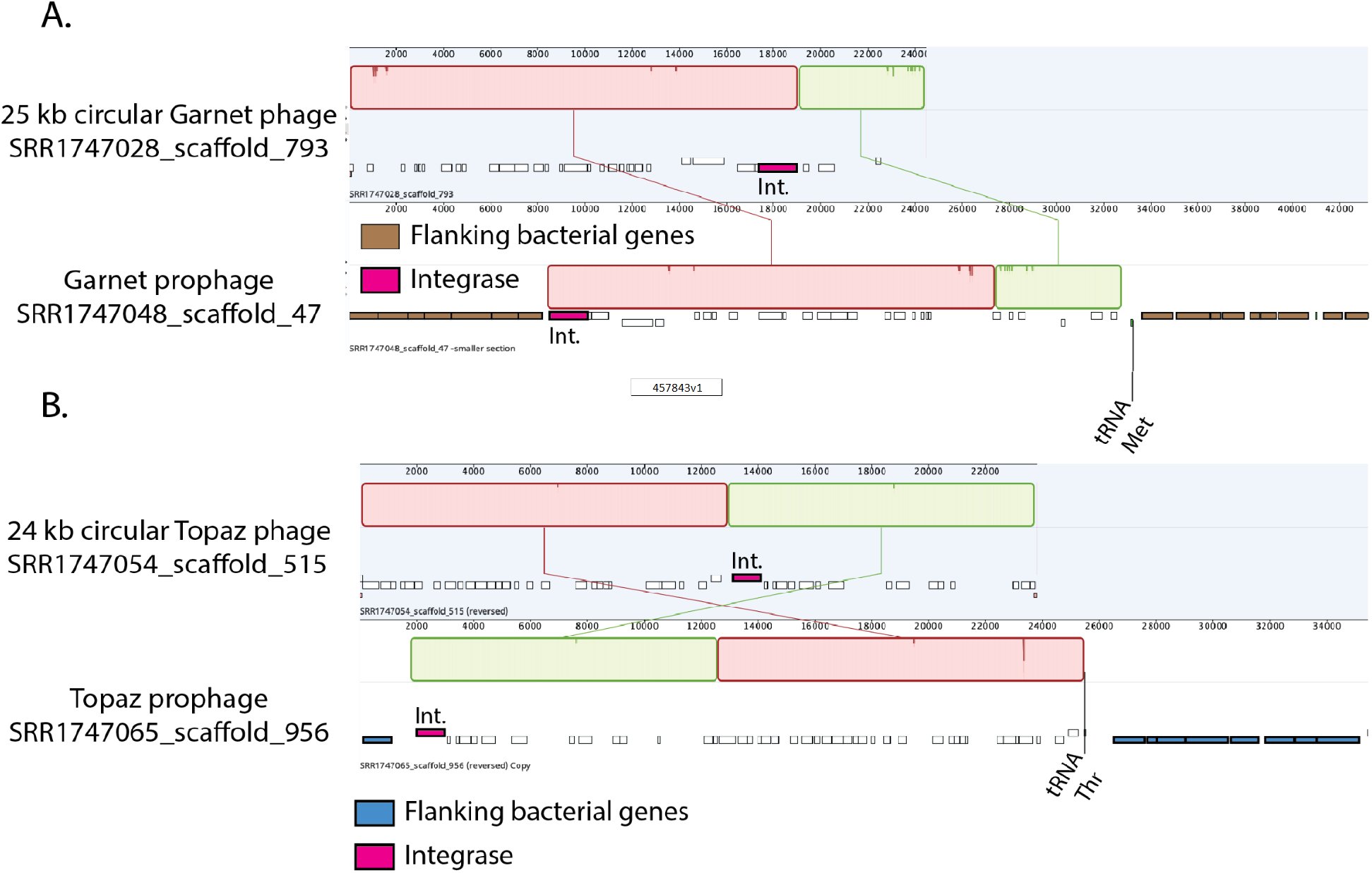
Full genome alignments between alternatively coded prophages and nearly-identical alternatively coded circular phages. **A**. A 25kb circular TAG-recoded Garnet phage aligned to a prophage integrated in a *Prevotella sp*. genome (*Prevotella* genes = brown). The prophage boundaries are marked by the phage integrase (pink) and the host tRNA Met. **B**. A 24kb circular TAG-recoded Topaz phage aligned to a prophage integrated into a *Oscillospiraceae sp*. genome (*Oscillospiraceae* genes = blue). The prophage boundaries are marked by the phage integrase (pink) and the host tRNA Thr.

## Supplementary Table Legends

**Table S1:** Clades of alternatively coded phages. Basic characteristics of the AC phage clades identified in this study. Circular genome size is shown as range from the smallest predicted circular genome in the clade to the largest predicted circular genome. checkV was used to evaluate genome completeness, and also identify artifactually long concatenated genomes.

| Name | Genetic Codes | Host Phylum | Circular Genome Size (kb) |
| --- | --- | --- | --- |
| Lak | TAG recoded | Bacteroidetes | 540-660 |
| crAss-like | Standard code<br>TAG recoded<br>TGA recoded | Bacteroidetes | 95-190 |
| Garnet | Standard code<br>TAG recoded | Bacteroidetes | 25-39 |
| Amethyst | Standard code<br>TAG recoded<br>TGA recoded | Bacteroidetes | 25-48 |
| Jade | TAG recoded<br>TGA recoded | Firmicutes | 150-224 |
| Sapphire | TAG recoded | Firmicutes | 209-213 |
| Agate | Standard code<br>TAG recoded<br>TGA recoded | Firmicutes | 106-201 |
| Topaz | Standard code<br>TAG recoded<br>TGA recoded | Firmicutes | 15-29 |

**Table S2:** Code change related genes in alternatively coded phages. A list of AC phage clades with release factors and tRNA synthetases. tRNAs are not shown for simplicity, as all AC phage clades had at least one genome with a predicted suppressor tRNA.

| Clade | Code | # Genomes with trait | Release Factor | tRNA synthetase |
| --- | --- | --- | --- | --- |
| Lak | TAG recoded | 6 | RF2 | – |
| Jade | TGA recoded | 2 | RF1 | Tryptophanyl tRNA synthetase |
| Agate | TAG recoded | 2 | RF2 | – |

